## Supplementary Information for "Basic reproduction number for pandemic *Escherichia coli* clones varies markedly and can be comparable to pandemic influenza viruses"

#### 1 Sensitivity analysis

As sensitivity analysis, we examined the impact of two parameters, relative invasiveness $\rho_c$ and the delay parameter $\Delta_t$ on the posterior distribution of $R_0$ and the fit of the model to data. Since the relative invasiveness (odds ratio) $\rho_c$ was an estimate itself, we chose to use the 95% confidence intervals for $\rho_c$ as sensitivity analysis: the model was fit for the lower 95% CI, mean value and upper 95% CI estimate. The mean value of $\rho_c$ was used for the main results.

In addition, we tested the impact of different priors for the delay parameter $\Delta_t$, which defines the delay between onset of colonization and infection. The prior is uniform in all cases, but three different maximum values for $\Delta_t$ were tested: 0 years (no delay), 0.5 years and 5 years. A maximum delay of 0.5 years was used in the main results.

In addition to the posterior distribution of $R_0$, we assess the sensitivity of our model to $\rho_c$ and $\Delta_t$ in terms of model fit. Supplementary Figures 2 and 3 show a visual comparison between different choices of $\rho_c$ and $\Delta_t$.

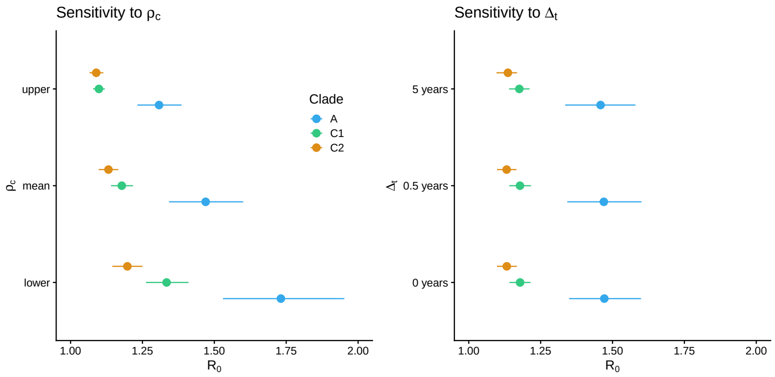


**Fig. 1** Impact of $\rho_c$ and $\Delta_t$ on the $R_0$ posterior distribution as a forest plot with 95% credible intervals as a horizontal line and the mean estimate as a point. On the left hand side, we display the effect of different $\rho_c$ estimates on the posterior distribution of $R_0$, colored by clade. On the right hand side, we display the 95% credible interval for $R_0$ for three different maximum delays: 0 years, 0.5 years and 5 years. Results for all clades seem identical regardless of delay value.

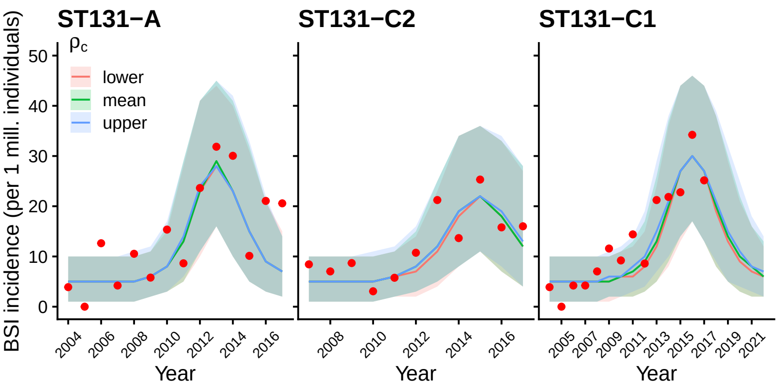


**Fig. 2** Effect of different $\rho_c$ estimates on the model fit to data. The different colors correspond to data fits using different $\rho_c$ estimates: lower and upper 95% CI and the mean $\rho_c$ estimate, which was used for the main results. The red dots correspond to observed data.

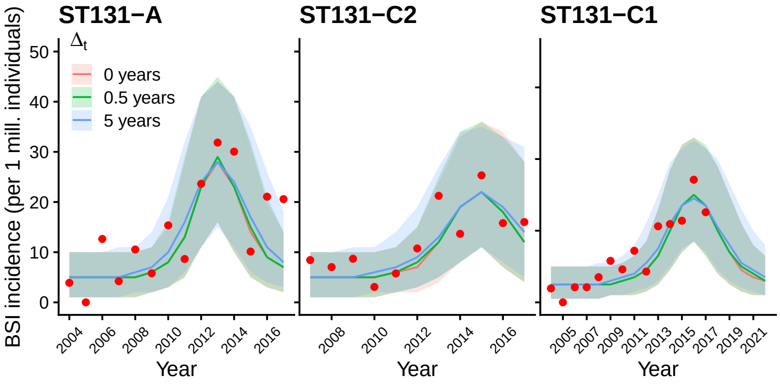


**Fig. 3** Effect of different maximum $\Delta_t$ values on the posterior predictive model fit to data. The red dots correspond to observed data. We can observe a slight increase in uncertainty as the maximum of the delay parameter is increased to 5 years.

### 2 Model comparison

We compare the chosen colonization model (SIR-based compartmental model), with an alternative SIS-based compartmental model. The SIS-based model does not assume a recovered/removed compartment and thus allows for re-colonization. For model comparison, we used ABC-SMC thresholds of 2.0, 1.5, 1.0 and 0.5 and locally optimal proposal distributions to allow for faster inference, since the SIS model was considerably more challenging to fit to the data. For C2 clade the simulation process was extended to 14 years due to difficulties of generating sufficiently many accepted particles from the posterior under the SIS model due to its poorer fit to data. For all clades, the acceptance rate of the SIS-based model is considerably lower for the smallest threshold of 0.5 than the acceptance rate for the SIR-based model. For ST131-C2, the decrease in acceptance rate starts is most notable, from threshold value of 1.0 to 0.5, as seen in Figure 5. From Figure 4 it can be seen that the fit to data is quite poor, especially for ST131-C2 and ST131-C1.

**Table 1** Acceptance rate (number of samples divided by number of simulations), number of samples, number of simulations and the threshold value for SIR and SIS models. Models were run for clades ST131-A, ST131-C2 and ST131-C1. Note that for ST131-A and ST131-C2 a locally optimal ABC-SMC algorithm was used.

| ST131 | SIR Acceptance rate | SIS Acceptance rate | Threshold |
|---|---|---|---|
| A | 0.1236 | 0.1643 | 2.0 |
| | 0.1360 | 0.0961 | 1.5 |
| | 0.0414 | 0.0444 | 1.0 |
| | 0.1204 | 0.0282 | 0.5 |
| C2 | 0.0943 | 0.0610 | 2.0 |
| | 0.0806 | 0.0506 | 1.5 |
| | 0.0146 | 0.0080 | 1.0 |
| | 0.0027 | 0.0001 | 0.5 |
| C1 | 0.0498 | 0.0358 | 2.0 |
| | 0.0350 | 0.0137 | 1.5 |
| | 0.0086 | 0.0018 | 1.0 |
| | 0.0071 | 0.0007 | 0.5 |

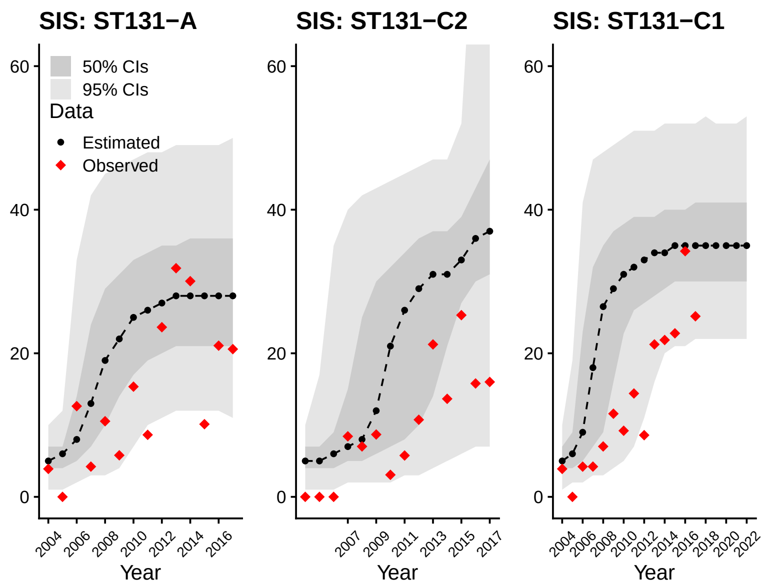


**Fig. 4** Fit of the SIS model to data. Red diamonds depict the observed data and black dots represent the mean of the estimated data. Grey and dark grey area are the 95 and 50 % credible intervals.

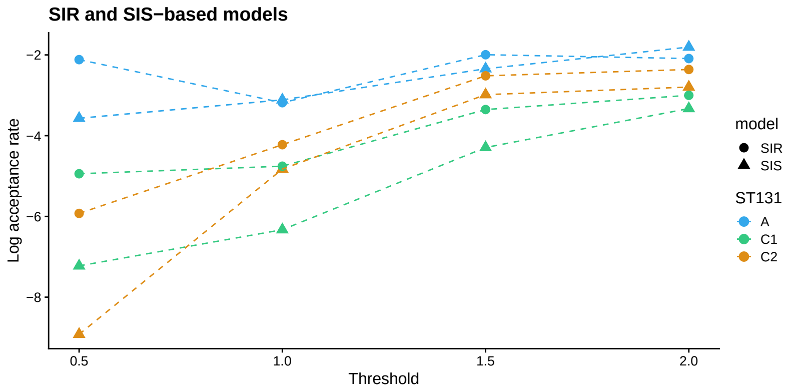


**Fig. 5** Logarithmic (natural logarithm) acceptance rate as a function of threshold for clades ST131-A, ST131-C2 and ST131-C1. Triangle-shaped points correspond to the SIS-based model and circular points to the SIR-based model.

### 3 Other Supplementary Items

#### 3.1 Identifiability of $R_0$, $\tau$ and $\Delta_t$

Plots of overlaid prior and posterior distributions of model parameters for clades ST131-A, ST131-C2 and ST131-C1 show that $R_0$ and $\tau$ are well identifiable, while the delay parameter $\Delta_t$ is not.

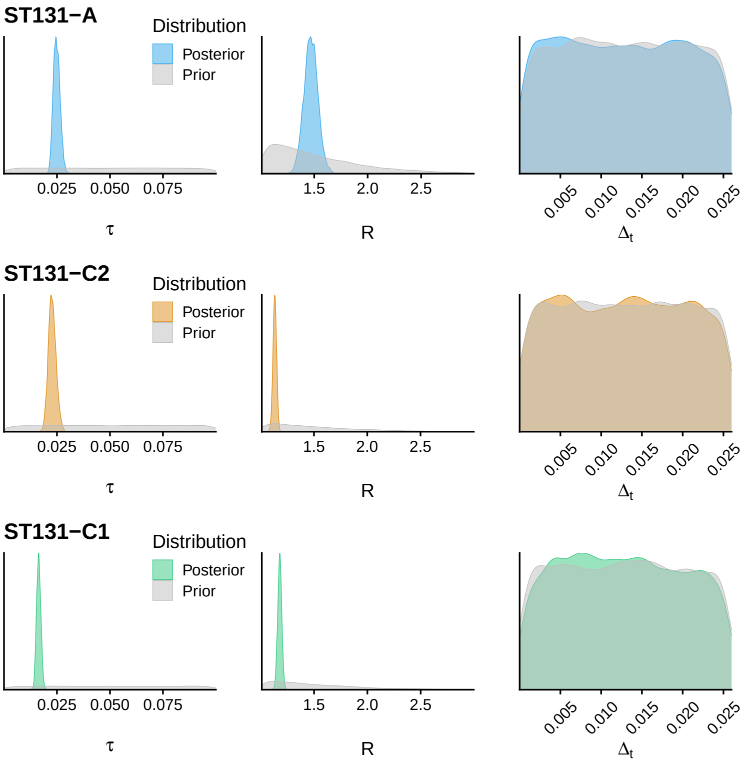


**Fig. 6** Overlay of the prior and posterior distributions for the estimated parameters $\tau$, $R_0$ and $\Delta_t$ for the clades of interest.

#### 3.2 Duration of colonization

In Figure 7, we show the posterior distribution of the duration of colonization for ST131-A, ST131-C2 and ST131-C1 in weeks. The duration of colonization is defined as $1/\gamma$, where $\gamma$ is the recovery rate of an SIR model. Refer to the main text for details on the model parametrization.

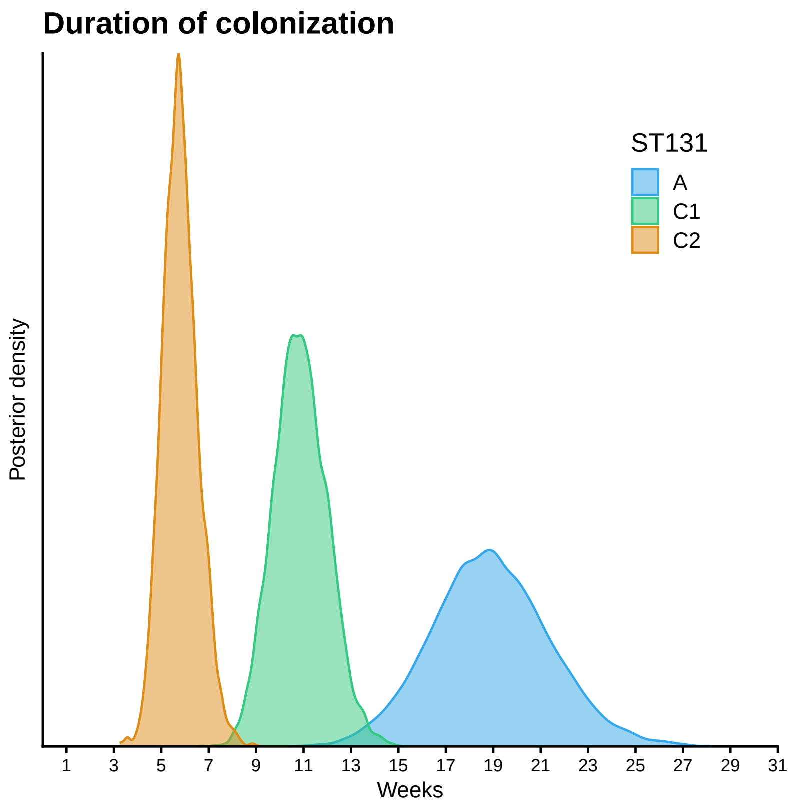


**Fig. 7** Posterior distribution of the duration of colonization in weeks, defined as $1/\gamma$, where $\gamma$ is the recovery rate of an SIR model.
